# Intranasal mRNA-LNP vaccination protects hamsters from SARS-CoV-2 infection

**DOI:** 10.1101/2023.01.11.523616

**Authors:** Gabriela Baldeon Vaca, Michelle Meyer, Ana Cadete, Chiaowen Joyce Hsiao, Anne Golding, Albert Jeon, Eric Jacquinet, Emily Azcue, Chenxia Monica Guan, Xavier Sanchez-Felix, Colette A. Pietzsch, Chad E. Mire, Matthew A. Hyde, Margaret E. Comeaux, Julie M. Williams, Jean C. Sung, Andrea Carfi, Darin K. Edwards, Alexander Bukreyev, Kapil Bahl

## Abstract

Intranasal vaccination represents a promising approach for preventing disease caused by respiratory pathogens by eliciting a mucosal immune response in the respiratory tract that may act as an early barrier to infection and transmission. This study investigated immunogenicity and protective efficacy of intranasally administered messenger RNA (mRNA)–lipid nanoparticle (LNP) encapsulated vaccines against severe acute respiratory syndrome coronavirus 2 (SARS-CoV-2) in Syrian golden hamsters. Intranasal mRNA-LNP vaccination systemically induced spike-specific binding (IgG and IgA) and neutralizing antibodies with similar robustness to intramuscular controls. Additionally, intranasal vaccination decreased viral loads in the respiratory tract, reduced lung pathology, and prevented weight loss after SARS-CoV-2 challenge. This is the first study to demonstrate successful immunogenicity and protection against respiratory viral infection by an intranasally administered mRNA-LNP vaccine.

## Introduction

Disease caused by respiratory pathogens remains a pre-eminent threat to global public health.^1^ With over 600 million cases and 6.5 million deaths reported worldwide as of November 2022, the ongoing coronavirus disease 2019 (COVID-19) pandemic caused by severe acute respiratory syndrome coronavirus 2 (SARS-CoV-2) is the most current and vivid example of the impact of respiratory diseases on global populations.^2^ Prior to the COVID-19 pandemic, upper and lower respiratory tract infections were responsible for over 17.7 billion cases and 2.5 million deaths globally, and primarily caused by viruses and bacteria such as *Streptococcus pneumoniae,* respiratory syncytial virus, and influenza virus.^3^ There remains a continual risk of emerging respiratory infectious diseases,^4^ as evidenced by evolving SARS-CoV-2 variants in the current COVID-19 pandemic as well as by notable prior pandemics caused by pathogens such as influenza virus.^2^ Vaccination remains a pivotal strategy to address infectious disease–related morbidity and mortality,^5^ with a need for innovative vaccination strategies and technologies that can be deployed quickly and establish robust local mucosal immune responses in the upper respiratory tract to impede infection and transmission.^6^

Currently, most licensed vaccines against respiratory diseases are administered intramuscularly, which primarily induce systemic immunity while also eliciting some immunity at the mucosal sites targeted by respiratory pathogens.^6–9^ Intranasal vaccination can induce both systemic and local mucosal immune responses, and is a promising approach to combat respiratory pathogens as it has the potential to limit infection and minimize transmission by establishing early, local immunity at key infection sites.^7,8,10,11,7,9,10,12–15^ This approach could also increase vaccination rates and compliance with recommended schedules, as its minimally invasive delivery may facilitate administration without the need for trained healthcare personnel.^7,16,17^ Additionally, intranasal vaccination by using a device to create a spray or aerosol could potentially bypass injection-associated phobias that are responsible for vaccine hesitancy.^18^ While few intranasal vaccines are currently authorized,^6,9^ the continued emergence of SARS-CoV-2 variants shifted attention to vaccination strategies that may better limit transmission and slow variant progression. Consequently, multiple intranasal SARS-CoV-2 vaccines based on viral vector, live attenuated, or protein subunit designs are currently in preclinical and clinical development,^7^ with 2 mucosal SARS-CoV-2 viral vectored vaccines having recently received regulatory approval in China and India.^19^

The messenger RNA (mRNA)–lipid nanoparticle (LNP) encapsulated vaccines have already demonstrated the ability to protect against infectious respiratory pathogens, as shown by currently available COVID-19 vaccines: mRNA-1273 (Spikevax; Moderna, Inc., Cambridge, MA, USA^20^) and BNT162b2 (Comirnaty; Pfizer Inc, New York, NY, USA; BioNTech Manufacturing GmbH, Mainz, Germany).^21–26^ Moreover, mRNA-LNP vaccines do not induce a vector-specific immune response and thus have high potential for repeat administration without diminishment of effect caused by anti-vector immunity.^27–29^ Further benefits include that mRNA is also non-infectious and non-integrative,^29^ while LNPs can be modified for delivery of mRNA to specific cells, tissues, and organs.^30,31^

In this report, we demonstrate that a 2-dose regimen of intranasally administered mRNA-based SARS-CoV-2 vaccines are immunogenic and protect against viral infection in a Syrian golden hamster model. These are the first known preclinical findings of an intranasally administered mRNA-LNP vaccine successfully protecting against infection by a respiratory pathogen.

## Results

### Intranasal mRNA-LNP vaccination induces binding and neutralizing antibody responses in sera

To assess the immunogenic potential of intranasally administered N1-methyl-pseudouridine-modified mRNA-LNPs, we developed SARS-CoV-2 vaccines formulated with 2 different LNP compositions: mRNA-LNP1 and mRNA-LNP2. mRNA-LNP1 is similar in composition to the LNP used in mRNA-1273, with similar but chemically distinct ionizable lipids, and mRNA-LNP2 is a composition further developed for improved respiratory tract delivery. All vaccines encoded a prefusion-stabilized SARS-CoV-2 spike (S) protein, stabilized with 6 proline mutations.^32^ Syrian golden hamsters (n = 10 per group) were vaccinated three weeks apart with 2 doses of either mRNA-LNP vaccines at 5 μg or 25 μg or with tris/sucrose buffer (mock-vaccinated) via the intranasal route (Days 0 and 21; **Figure 1**). For comparison purposes, 2 groups of hamsters were intramuscularly vaccinated with 0.4 μg or 1 μg of vaccine with the same mRNA included in the intranasal compositions but formulated with the preclinical version of the same LNP utilized in injectable mRNA-1273. Immunogenicity was assessed at approximately 3 weeks after dose 1 (Day 21) and approximately 3 weeks after dose 2 (Day 41); S-specific serum immunoglobulin (Ig) G or A binding antibody responses were measured by enzyme-linked immunosorbent assay (ELISA) and serum neutralizing antibody titers were measured by a plaque reduction neutralization test (PRNT).

**Figure 1.**
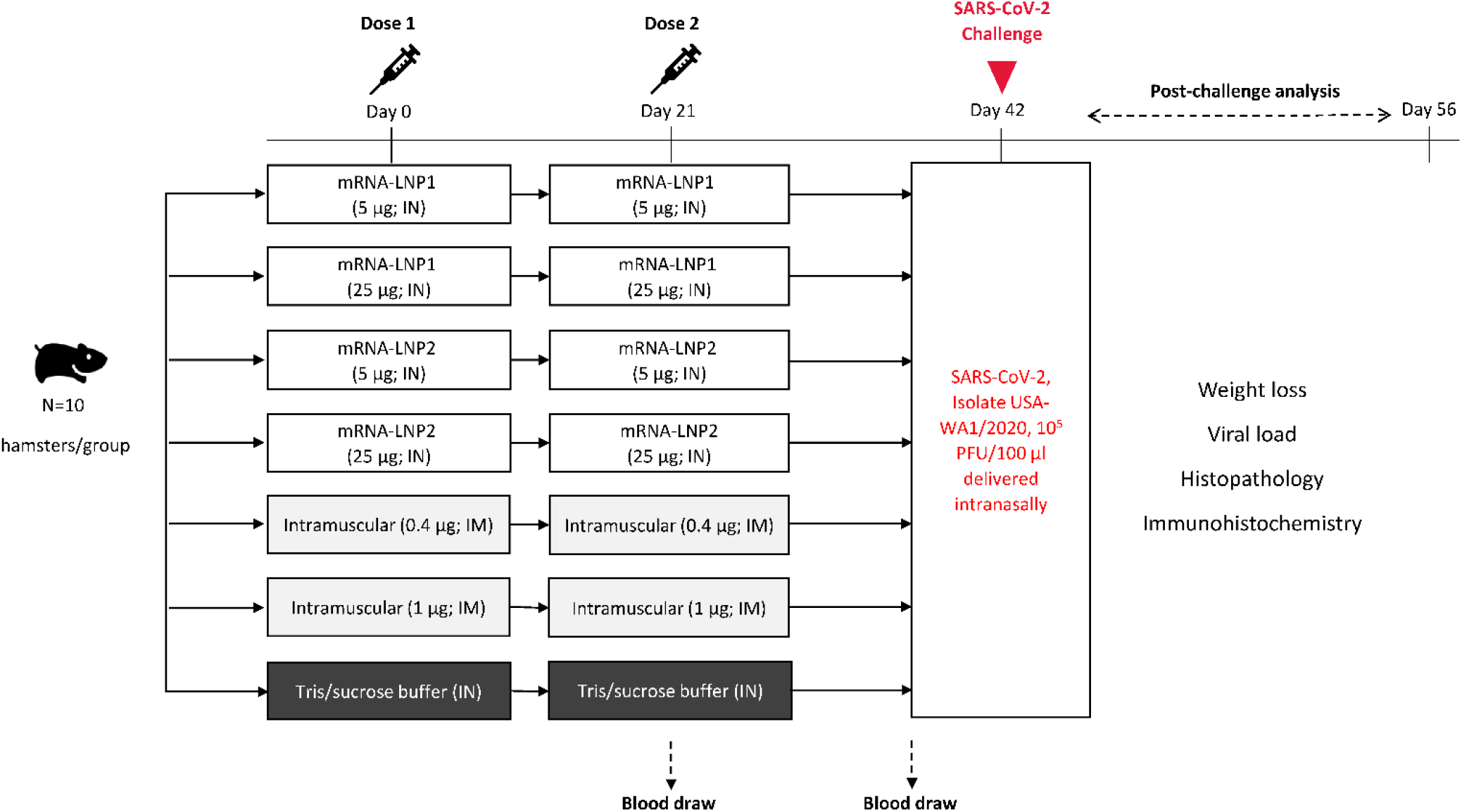
Study design. Intranasal vaccination of an mRNA-based SARS-CoV-2 vaccine was evaluated in Syrian golden hamsters. Hamsters (n = 10 per group) were intranasally immunized with 2 doses (Day 0 and Day 21) of vaccines (5 μg or 25 μg) formulated in 2 different LNP compositions or were mock-vaccinated with 2 doses of tris/sucrose buffer administered intranasally; separate groups of animals were intramuscularly immunized with 2 doses of vaccine (0.4 μg or 1 μg). Sera were collected approximately 3 weeks after dose 1 (prior to dose 2 on Day 21) and approximately 3 weeks after dose 2 (Day 41). At Day 42, hamsters were intranasally challenged with SARS-CoV-2 (2019-nCOV/USA-WA1/2020). Post–viral challenge assessments included viral load and histopathology (3 days [Day 45] and 14 days [Day 56] after challenge), immunohistochemistry (3 and 14 days after challenge), as well as body weight (daily after challenge). IM, intramuscular; IN, intranasal; LNP, lipid nanoparticle; mRNA, messenger RNA; PFU, plaque-forming units; SARS-CoV-2, severe acute respiratory syndrome coronavirus 2.

Three weeks after the first dose, both intranasal vaccines (25-μg dose level) elicited S-specific serum IgG binding titers comparable to those induced by intramuscular administration (0.4 μg and 1 μg). At the 5-μg dose level, mRNA-LNP2 induced similar titers to mRNA-LNP2 25 μg and to intramuscular controls (0.4 μg and 1 μg); mRNA-LNP2 titers at the lower 5-μg dose were significantly higher than mRNA-LNP1 titers (adjusted *P*<0.0001; **Figure 2a**; **Table S1**). After the second dose, S-specific IgG titers generally increased across all vaccine groups and dose levels, with mRNA-LNP2 eliciting significantly higher titers than mRNA-LNP1 at the corresponding dose levels (5 μg, adjusted *P*<0.01; 25 μg, adjusted *P*<0.001).

**Figure 2.**
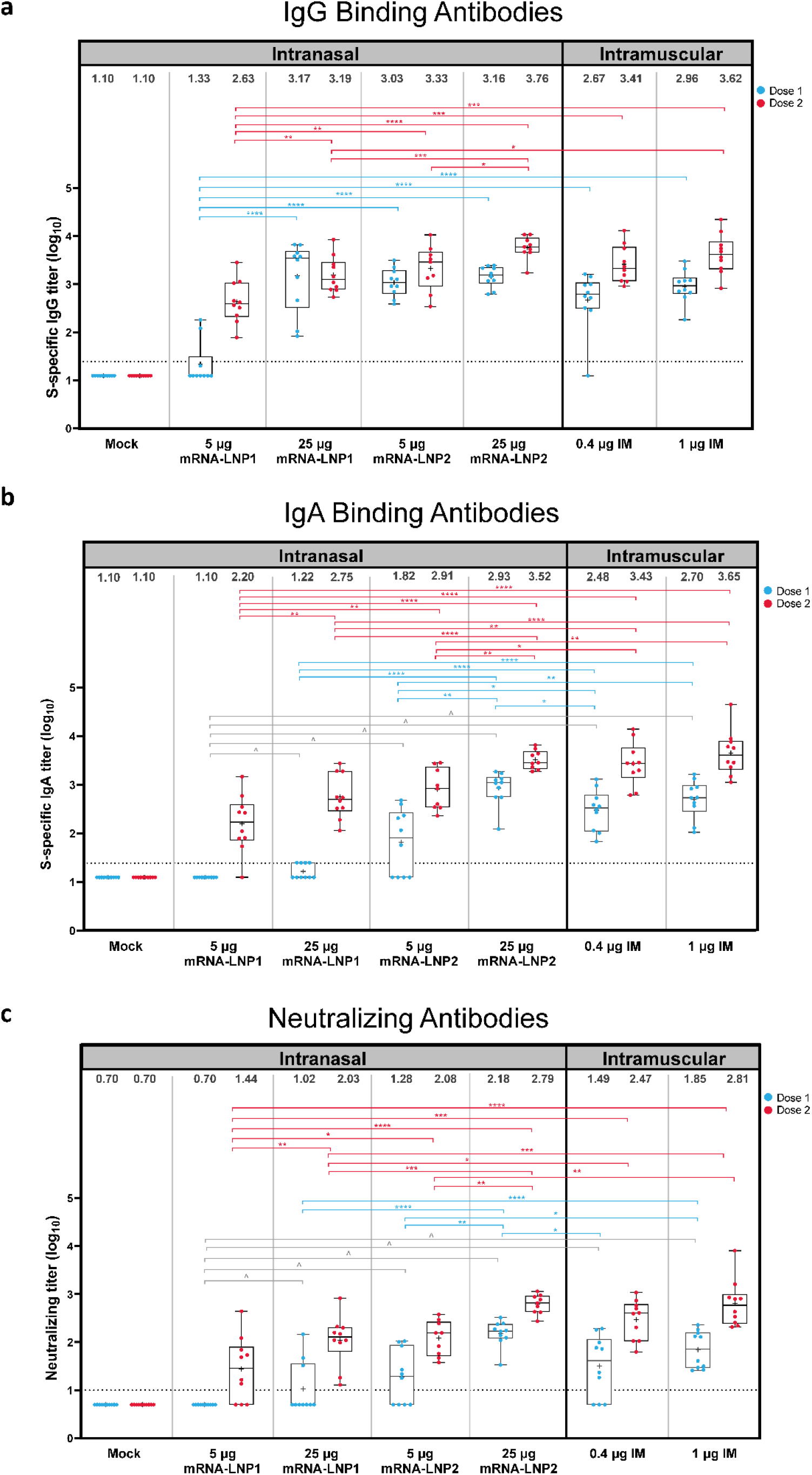
Serum immune responses after intranasal vaccination. (a) S-specific serum binding IgG, (b) S-specific serum binding IgA, and (c) serum neutralizing antibody reciprocal endpoint titers (log scale) at 3 weeks after dose 1 (Day 21) or 3 weeks after dose 2 (Day 41) are shown by vaccine group. Animal-level data are shown as dots (n = 9-10 animals per group), with boxes and horizontal bars denoting the IQR and median, respectively, and whiskers representing the maximum and minimum values. Geometric mean titers for each vaccine group are indicated by the plus (+) symbol of each boxplot, with the exact values shown above each vaccine group. Horizontal dotted lines represent the LLOD. *P<.05, **P<.01, ***P<.001, ****P<.0001. Results of statistical comparisons between groups are shown in **Tables S1-S3**. ^Antibodies were under the limit of detection for all hamsters in the mRNA-LNP1 5 μg group after dose 1, which had a much lower antibody level compared to other groups. IgA, immunoglobulin A; IgG, immunoglobulin G; IM, intramuscular; IN, intranasal; LLOD, lower limit of detection; LNP, lipid nanoparticle; mRNA, messenger RNA; S2-P, S-protein with 2 proline mutations; SD, standard deviation.

A single intranasal administration of mRNA-LNP2 (5 μg and 25 μg) elicited S-specific serum IgA binding antibody titers in sera (**Figure 2b**), with the 25-μg dose level eliciting higher (adjusted *P*<0.05) or similar titers as intramuscular administration (0.4 μg and 1 μg, respectively; **Table S2**). mRNA-LNP1 at the 5-μg and 25-μg dose levels elicited lower titers than intramuscular controls and respective mRNA-LNP2 doses. A second dose of either intranasal vaccine composition increased IgA binding titers, with the 25-μg dose levels eliciting significantly higher titers than the respective 5-μg-dose level (adjusted *P*<0.01). Additionally, mRNA-LNP2 25 μg elicited higher or comparable IgA titers to intramuscular administration (0.4 μg and 1 μg) after either dose.

In addition to S-specific binding titers, neutralizing antibody responses in sera were evaluated (**Figure 2c**). Three weeks after the first dose, titers were not detected in some hamsters after mRNA-LNP1 (25 μg dose), mRNA-LNP2 (5 μg dose), or intramuscular administration (0.4 μg dose). No hamsters administered mRNA-LNP1 5 μg had detectable titers after the first dose. However, all hamsters vaccinated with mRNA-LNP2 25 μg had neutralizing antibody titers that were significantly higher (*P*<0.05) or comparable to intramuscular controls (0.4 μg and 1 μg, respectively; **Table S3**). Neutralizing antibody titers increased for all vaccine groups after the second dose. mRNA-LNP2 (5 μg and 25 μg) induced significantly higher titers than mRNA-LNP1 at the respective dose level (5 μg, adjusted *P*<0.05; 25 μg, adjusted *P*<0.001). Two doses of mRNA-LNP2 25 μg induced similar neutralizing titers to intramuscular vaccination (0.4 μg and 1 μg).

### Intranasal mRNA-LNP vaccination limits viral replication in the respiratory tract and protects against disease

Three weeks after the second dose (Day 42), all vaccinated and mock-vaccinated hamsters were challenged intranasally with 10^5^ plaque-forming units (PFU) of SARS-CoV-2 (isolate USA-WA1/2020; **Figure 1**). This isolate was selected for challenge as ancestral SARS-CoV-2 isolates are more pathogenic and drive more severe disease in hamsters than omicron lineage viruses.^33,34^ Viral loads in nasal turbinates and lung were then assessed 3 days (Day 45; n = 5 animals per group) and 14 days after challenge (Day 56; n = 5 animals per group) and body weight was evaluated daily.

At 3 days after SARS-CoV-2 challenge, intranasally vaccinated hamsters had lower viral loads in both the lung and nasal turbinates relative to mock-vaccination, as determined by plaque assay (**Figure 3a** and **3b**, respectively). In the lung, viral loads were below the levels of detection in 4 of 5 hamsters vaccinated with mRNA-LNP2 25 μg, which was significantly reduced relative mock-vaccinated controls (*P*<0.05; **Table S4**). Viral loads were not detected in 2 of 5 hamsters vaccinated with mRNA-LNP2 5 μg or mRNA-LNP1 25 μg, similar to the 0.4 μg intramuscular vaccine dose level (2 of 5 animals). At the 1 μg intramuscular dose level, which is considered protective in this hamster model,^35^ viral load in the lungs was significantly reduced compared with mock-vaccinated controls (*P*<0.05), with 3 of 5 vaccinated hamsters having no detectable virus. Overall, viral loads in the lung were lower among hamsters intranasally administered mRNA-LNP2 than mRNA-LNP1 at the respective dose levels, which was significant at the 5-μg dose level (*P*<0.05). Similarly, in nasal turbinates, viral loads were undetected in 1 of 5 hamsters vaccinated with mRNA-LNP1 25 μg and 2 of 5 hamsters vaccinated with mRNA-LNP2 25 μg; loads were significantly lower with mRNA-LNP2 25 μg than mock-vaccination (*P*<0.01; **Table S5**). Viral titers among hamsters intranasally vaccinated with the 5 μg-dose level of either intranasal composition remained detectable at 3 days after infection, but titers were numerically lower relative to mock-vaccination and were generally similar to intramuscular vaccination (0.4 μg). Overall, viral reduction in the lungs and nasal turbinates of intranasally vaccinated groups was comparable to intramuscularly vaccinated groups at the respective lower or higher dose level. By 14 days after challenge, SARS-CoV-2 virus was not detectable in lung or nasal turbinates of any intranasally or intramuscularly vaccinated hamsters, including mock-vaccinated animals (**Figure 3a** and **3b**).

**Figure 3.**
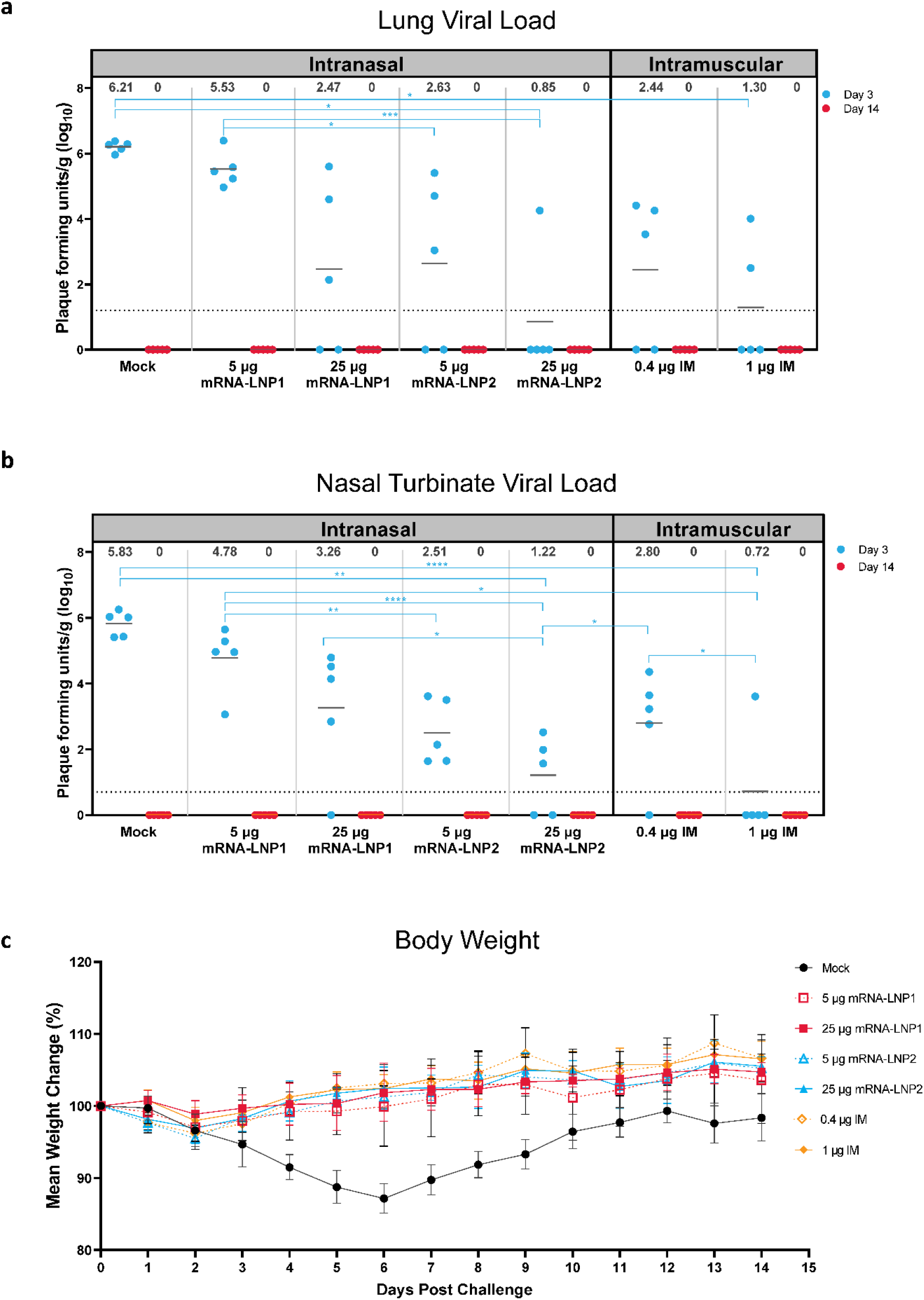
Viral load and weight loss characteristics after SARS-CoV-2 challenge in vaccinated hamsters. (a) Viral load (PFU per gram of tissue) in lungs and (b) viral load in nasal turbinates of mock-vaccinated and vaccinated hamsters at 3 days and 14 days after SARS-CoV-2 challenge. Animal-level data are shown as dots (n = 5 animals per group), with grey lines representing the geometric mean titer for each group; exact values are shown above each vaccine group. Statistical comparisons were only performed for viral loads at day 3 after challenge, as viral loads at day 14 were zero for all hamsters. *P<.05, **P<.01, ***P<.001, ****P<.0001. Results of statistical comparisons between groups are shown in **Tables S4-S5**. (c) Mean percentage of weight change (error bars represent SEM) over 14 days after SARS-CoV-2 challenge in mock-vaccinated and vaccinated hamsters. IM, intramuscular; IN, intranasal; LNP, lipid nanoparticle; mRNA, messenger RNA; PFU, plaque-forming units; SARS-CoV-2, severe acute respiratory syndrome coronavirus 2; SEM, standard error of the mean.

Viral load in respiratory tissues was also determined through assessment of viral subgenomic RNA (sgRNA) levels by quantitative reverse transcription polymerase chain reaction (qRT-PCR). Corroborating the plaque assay results, all vaccinated hamsters at 3 days post SARS-CoV-2 challenge had slightly lower viral sgRNA levels relative to mock-vaccinated controls, regardless of dosage and route of administration (**Figure S1**). By 14 days after challenge, sgRNA was not detectable in lung or nasal turbinates of any hamsters, including those mock vaccinated.

SARS-CoV-2 infection was performed with a sub-lethal dose known to result in disease characteristics such as weight-loss in Syrian golden hamsters.^36^ Over the course of infection, mock-vaccinated hamsters experienced a maximum mean (± standard error) weight loss of 12.9% (± 1.02) by day 6 post-challenge (**Figure 3c**). Comparatively, all intranasally or intramuscularly vaccinated hamsters maintained their bodyweights over the 14-day post-challenge period.

### Intranasal mRNA-LNP vaccination reduces severity of viral pathology in the lungs

In the Syrian golden hamster model, SARS-CoV-2 infection with ancestral strains causes severe pathological lesions in lung tissue by 3 days after infection that typically begins to resolve by 10 days after infection.^36^ Therefore, to examine the ability of intranasal mRNA-LNP vaccination to reduce lung pathology after infection, histopathological examination of the lower left lobe of the lung of hamsters was performed at 3 days and 14 days after challenge.

Three days after infection, all vaccinated and mock-vaccinated hamsters exhibited acute pulmonary parenchymal tissue damage and inflammation. There were regionally extensive areas of interstitial infiltration by mixed inflammatory cells, alveolar accumulation of fibrin, hemorrhage, infiltration of bronchial/bronchiolar epithelium by neutrophils, large clusters of intraluminal neutrophils within bronchi/bronchioles with or without epithelial degeneration/necrosis, and vascular inflammation. However, there were vaccine group- and dose-dependent differences in severity. Hamsters vaccinated with high dose levels of either intranasal (25 μg) or intramuscular (1 μg) vaccine compositions had similar levels of pulmonary parenchymal inflammation (**Figure 4a**) to mock-vaccinated controls but exhibited lower severity scores for bronchial/bronchiolar inflammation (**Figure 4b**) and vascular inflammation (**Figure 4c**). No major differences in lung histopathology were observed for the lower dose levels compared to mock-vaccination (**Table S6**). Major histopathological findings and severity scores for each group and dose level at 3 days after challenge are summarized in **Table S6**.

**Figure 4.**
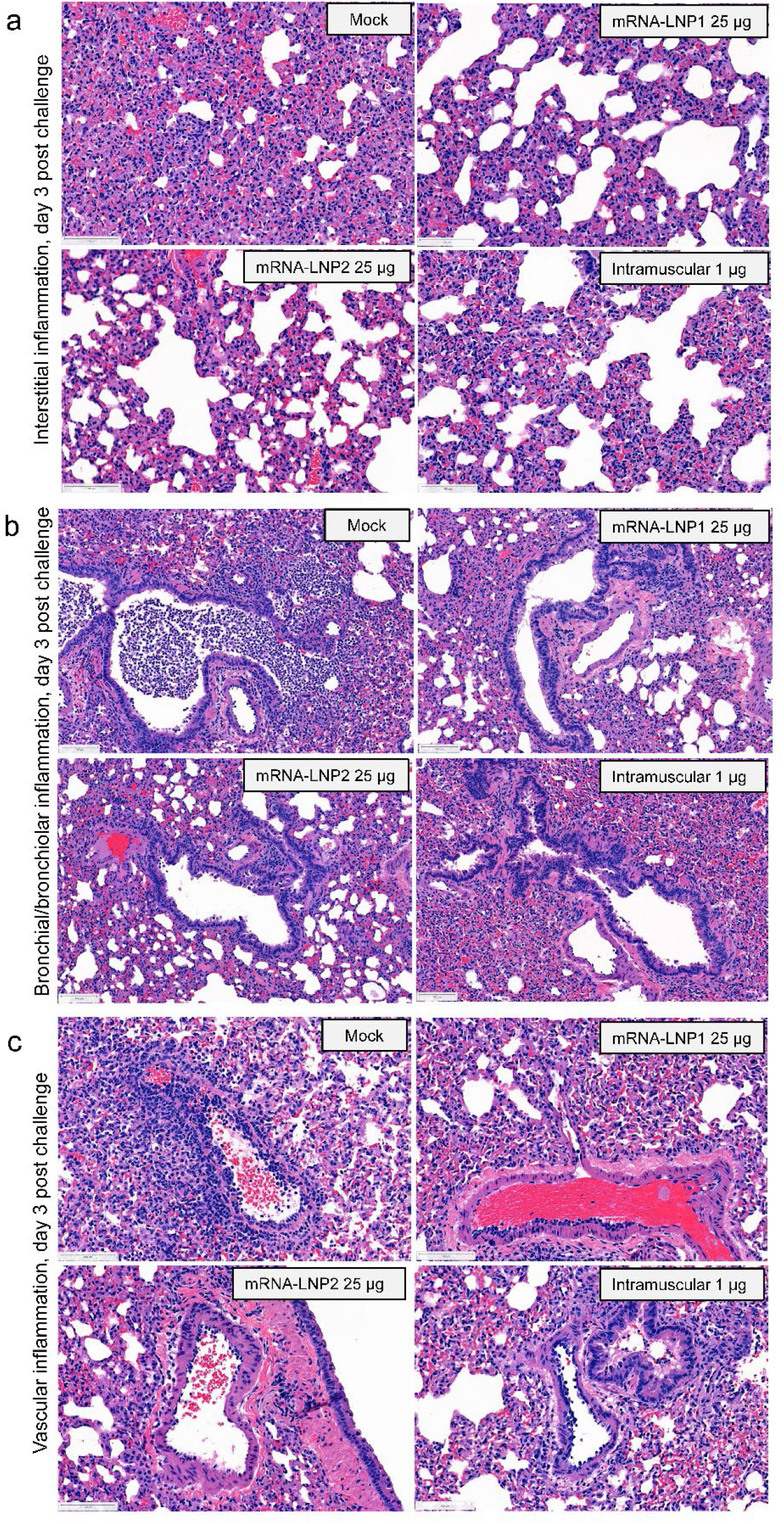
Pulmonary histopathological characteristics at 3 days after SARS-CoV-2 challenge in vaccinated hamsters. Lung sections from hamsters at 3 days after SARS-CoV-2 challenge were stained with H&E. Representative images are shown for mock-vaccinated, intranasally vaccinated (25 μg), or intramuscularly immunized (1 μg) hamsters. (a) The pulmonary parenchyma show moderate, interstitial infiltration by mixed inflammatory cells within alveolar walls, multifocal deposits of fibrin, and alveolar hemorrhage. (b) Airways including bronchi and bronchioles were frequently obstructed by high numbers of neutrophils in mock-vaccinated hamsters. Note the lack of this suppurative inflammation in vaccinated hamsters. (c) Vascular and perivascular mixed cell infiltrates were observed in medium to large-sized blood vessels. Note the decreased severity of vascular inflammation in vaccinated hamsters. Scale bars represent 100 μm. H&E, hematoxylin and eosin; IN, intranasal; LNP, lipid nanoparticle; mRNA, messenger RNA; SARS-CoV-2, severe acute respiratory syndrome coronavirus 2.

Fourteen days after SARS-CoV-2 infection, there were still regionally extensive areas of interstitial inflammation for all hamsters regardless of administration route or dose level (**Table S6**). However, fibrin accumulation, hemorrhage, bronchial/bronchiolar inflammation, and vascular inflammation regressed, with evidence of tissue recovery such as type II pneumocyte hyperplasia. Nonetheless, compared with mock-vaccination, all vaccinated groups exhibited lower severity of pulmonary inflammation irrespective of vaccine group or dose level (**Table S6**). Histopathology at 14 days after challenge for high dose levels are shown in **Figure S2**.

Additionally, lung tissue samples were stained for the SARS-CoV-2 nucleocapsid protein (N protein) by immunohistochemistry to identify cells infected with SARS-CoV-2 (**Figure 5a**). Three days after challenge, all 5 mock-vaccinated hamsters had N-protein+ cells (group mean ± standard error of the mean [SEM]: 43.5% ± 8.6% positive cells of total cells quantified; **Figure 5b**; **Table S7**). While N-protein was detected in the lung tissue of some vaccinated hamsters, vaccinated groups had a lower percentage of N-protein+ cells compared to mock-vaccinated controls (mRNA-LNP1: 5 μg, 8.4 ± 3.1%; 25 μg, 1.0 ± 0.7%; mRNA-LNP2: 5 μg, 6.4 % ± 3.8; 25 μg, 4.5% ± 4.5%; IM: 0.4 μg, 1.5% ± 0.8; 1.0 μg, 1.7% ± 1.2). The reduced percentage of N-protein+ cells relative to mock-vaccinated controls was significant for higher dose level groups, regardless of administration route (mRNA-LNP1 25 μg, *P*=0.017; mRNA-LNP2 25 μg, *P*=0.033; intramuscular 1 μg, *P*=0.004). Percentage of N-protein+ cells for lower dose level groups were not significantly different from mock-vaccinated controls. Of note, the mRNA-LNP2 25 μg group had 4 of 5 hamsters with <1% of N-protein positive cells, with one hamster having 22.6% N-protein+ cells. Relative to mock-vaccination, both low and high dose levels of intranasal vaccines reduced the percentage of N-protein+ cells in the lung, while the higher dose levels better ameliorated the pathologic manifestation. By 14 days after challenge, no groups were positive for N-protein, indicating virus clearance from the lungs (**Figure 5a** and **Figure 5b**).

**Figure 5.**
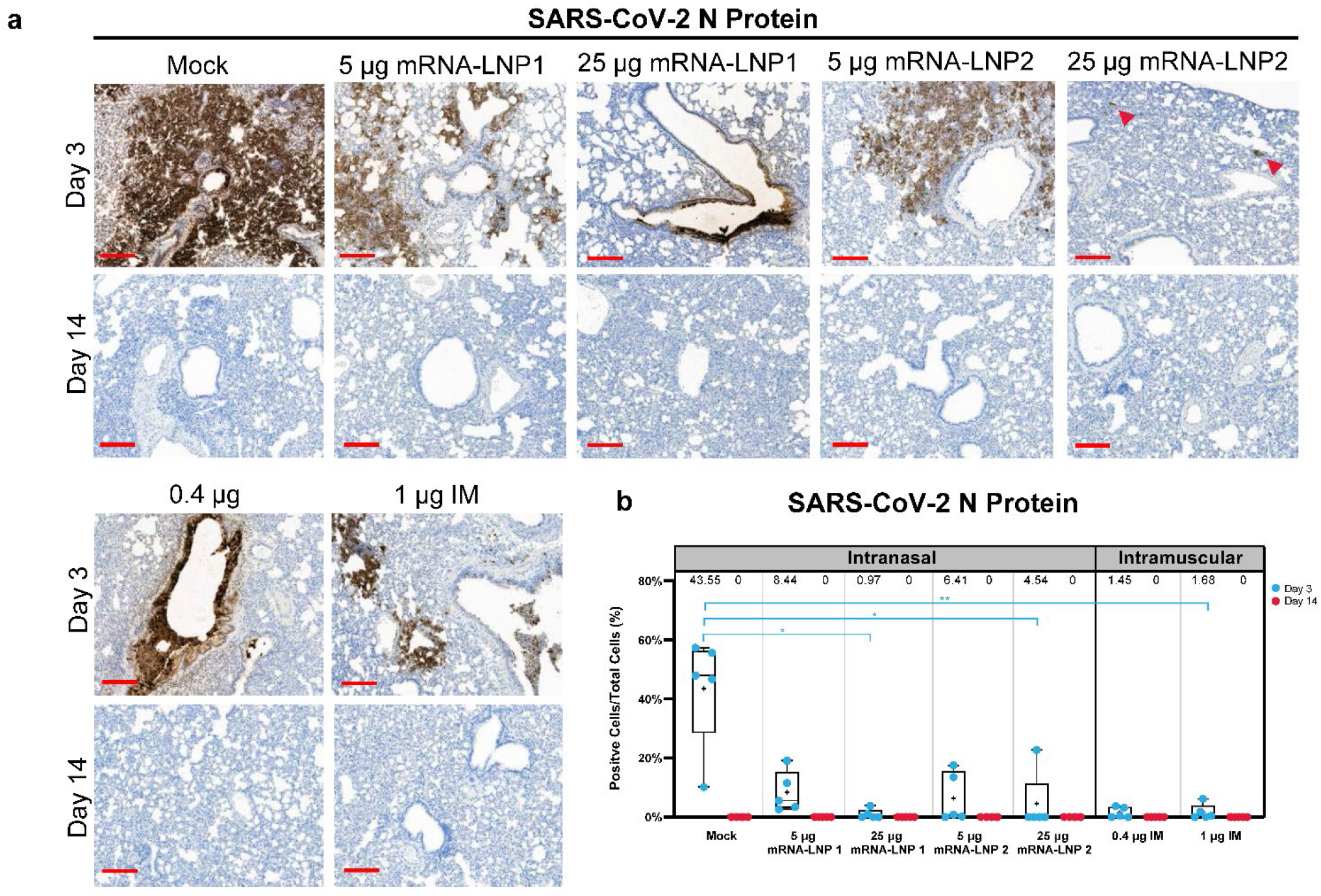
Immunohistochemistry for SARS-CoV-2 N-protein in lung after SARS-CoV-2 challenge. Lung sections from hamsters necropsied at 3 and 14 days after SARS-CoV-2 challenge were stained with an antibody raised against the SARS-CoV-2 N-protein. (a) Representational images of lungs from mock-vaccinated, intranasally vaccinated (mRNA-LNP1 or mRNA-LNP2 [5 μg and 25 μg]), or intramuscularly vaccinated (0.4 μg and 1 μg) hamsters. Arrowheads designate areas of positive signal within tissue. (b) Quantification of N-protein+ cells by vaccine group. Animal-level data are shown as dots (n = 4-5 animals per group), with boxes and horizontal bars denoting the IQR and median, respectively, and whiskers representing the maximum and minimum values. Mean values are provided above each plot. Scale bars represent 200 μm. N = 5 animals per group. Kruskal-Wallis non-parametric test was implemented for statistical analysis to accommodate for small sample sizes per group and subsequent small percentage of N-protein+ cells within vaccinated animals.\**P*< 0.05 and \*\**P* < 0.01. IN, intranasal; IQR, interquartile range; LNP, lipid nanoparticle; mRNA, messenger RNA; N protein, nucleocapsid protein; SARS-CoV-2, severe acute respiratory syndrome coronavirus 2.

## Discussion

Intranasal vaccine regimens may establish local immunity at upper respiratory sites and act as an early, protective barrier to reduce viral infection and subsequent transmission.^7,8,10,11^ However, vaccine development for intranasal administration is challenging. The respiratory tract is protected by a slightly acidic mucosal layer containing proteolytic enzymes that form a barrier over the epithelial cell lining that undergoes continuous mucosal clearance.^8^ These mechanisms act to defend against entry of respiratory pathogens but can subsequently prohibit antigen delivery during intranasal vaccination.^8^ Thus, novel technologies are needed to overcome these physiological barriers to advance intranasal vaccination and protect against respiratory disease.

Multiple intranasal vaccines against SARS-CoV-2 (using adenoviral vectors, live attenuated recombinant viruses, or adjuvant-protein subunits) are under investigation in both preclinical and clinical studies.^7^ Further, 2 viral-vectored mucosal vaccines were recently approved as booster doses in India (iNCOVACC; intranasal delivery through nasal drops; Bharat Biotech International Limited; Hyderabad, India)^37^ and China (Convidecia Air; nebulized vaccine for inhalation through the mouth; CanSino Biologics Inc.; Tianjin, People’s Republic of China).^38^ Collectively, preclinical studies examining intranasal SARS-CoV-2 vaccines have demonstrated induction of both systemic and local antibody responses and subsequent protection against SARS-CoV-2 infection,^10,15,39–41^ providing overall support for this vaccination route. However, findings from a recent phase 1 clinical trial on an adenovirus-vectored COVID-19 vaccine (ChAdOx1 nCoV-19) highlight the difficulties of translating animal research to humans, as the intranasal vaccine did not consistently elicit robust mucosal or systemic immune responses in vaccine-naive or previously vaccinated participants^42^ despite promising preclinical data.^15^

An mRNA-LNP–based approach for intranasal vaccination to respiratory pathogens, including SARS-CoV-2, may offer additional advantages over more traditional vaccine development platforms.^7,17^ For example, mRNA-encoded antigens more closely resemble the structure and presentation of viral proteins expressed during a natural infection.^43^ Additionally, an mRNA-based approach uses a single vaccine platform across different pathogens,^43^ with this platform enabling flexible antigen design, inclusion of multiple or modified antigens, and rapid incorporation of sequence substitutions that may be needed due to the emergence of variants.^43^ mRNA-based vaccines may also potentially minimize safety concerns associated with more traditional approaches utilized for mucosal vaccines, including those reliant on a live attenuated virus that have a theoretical risk of reverting to its pathogenic form. In addition, mRNA vaccines have a vector-less approach and thus can avoid the potential for diminished immunogenicity with repeat dosing sometimes observed with vector-based vaccines. Moreover, the utility of intramuscularly administered mRNA vaccines against respiratory pathogens such as SARS-CoV-2 has been established, demonstrating robust immune responses and high real-world effectiveness against disease.^22,44,45^ However, adapting this platform for intranasal vaccination still poses known technical challenges, including identifying the key target cells in the respiratory tract and as well as sufficient mucoadhesion and penetration to access these cells. Additional unknown hurdles to developing an immunogenic and effective mRNA-LNP intranasal vaccine may also be uncovered as this burgeoning approach continues to be investigated and advanced in the field.

This preclinical study explored the immunogenicity and protective efficacy of intranasally administered mRNA-LNP vaccines using SARS-CoV-2 as a model pathogen. Overall, a 2-dose primary intranasal vaccination regimen elicited systemic immune responses and resulted in lower SARS-CoV-2 infection levels and disease severity versus mock-vaccinated controls after viral challenge. In particular, 2 doses of mRNA-LNP2 elicited systemic immune responses that were generally similar to intramuscular administration (0.4 μg and 1.0 μg). Further, vaccination with mRNA-LNP2 reduced post-challenge viral titers in the lung and nasal turbinates relative to mRNA-LNP1 at the respective 5-μg and 25-μg dose levels, suggesting improved protection against SARS-CoV-2. Both intranasal vaccine formulations at the 25-μg level prevented severe lung pathology and reduced SARS-CoV-2 infection within the lungs to a similar degree as intramuscular vaccination. Taken together, these findings indicate that intranasal vaccination with an mRNA-LNP SARS-CoV-2 vaccine is protective and can induce systemic immune responses similar to intramuscular vaccination, which has already been shown to be highly effective against COVID-19.^21,22,46^

The findings of this study should be considered alongside several limitations. First, S-specific mucosal IgA levels were not specifically measured due to both bronchoalveolar lavage and nasal wash procedures being terminal in hamsters; therefore, mucosal-specific antibody responses resulting from intranasal vaccination were not determined in this study. However, it would be expected that intranasal vaccination would elicit higher mucosal IgA immune responses in the respiratory tract than intramuscular vaccination.^39,41^ Our finding that intranasal vaccination with the higher dose level of mRNA-LNP2 induced comparable serum binding IgA titers to an intramuscular route remains encouraging. A further limitation to this study is that efficacy assessments were performed in a preclinical animal model that is known to be highly susceptible to SARS-CoV-2 infection,^36,47^ which may not be immediately translatable to other animal models or human populations.

Additional studies that further evaluate the potential advantages of intranasal vaccination in preclinical models other than an acute protection model should assist in translating these findings to clinical settings. Studies in ferrets (in addition to hamsters) can aid in examining the potential for intranasal vaccination to reduce transmission, while studies in mice and non-human primate models could enable investigating persistence of immune responses and the induction of local mucosal tissue resident immunity and cellular immunity. However, the technically challenging nature of intranasal vaccination, coupled with the limited predictive power of preclinical intranasal vaccine findings for human populations, will need to be considered throughout vaccine development. Intranasal vaccination as a booster regimen following primary parenteral vaccination schedules should also be evaluated, as an intranasal booster could build upon primary vaccination to supplement mucosal immunity and provide early, durable protection against infection. Notably, the selected intranasal vaccine dose levels investigated in this study were exploratory and based on an unpublished pilot study in mice focused on protein expression; intramuscular dose levels were based on a previous study in hamsters.^35^ Use of an intranasal spray or an aerosolization device could possibly benefit vaccine delivery by allowing for improved delivery throughout the respiratory tract.

In conclusion, we have demonstrated that intranasally administered mRNA-LNP vaccines delivered as a primary 2-dose regimen are immunogenic and can protect naive hamsters from SARS-CoV-2 infection. Further, LNPs designed for improved delivery to the respiratory tract were more immunogenic and better protected against infection than traditional LNPs delivered intranasally. These encouraging findings may have broad implications for a progressive vaccination approach to supplement or complement current intramuscular approaches. Further investigations of intranasal mRNA-LNP vaccination alone or as a booster dose following an intramuscular primary series are thus warranted to address the burden of infectious respiratory disease worldwide. While an mRNA-based approach to intranasal vaccination may need to be further developed to reduce the effective dose level and increase suitability for human populations, our promising results show that mRNA-LNP vaccines have potential for effective administration via the intranasal route.

## Methods

### Hamster studies

Female Syrian golden hamsters (6-7 weeks old; Envigo) were intranasally vaccinated with a SARS-CoV-2 vaccine on a 2-dose schedule with 3 weeks between doses (Day 0 and Day 21; **Figure 1**). Hamsters (n = 10 per vaccine group) were intranasally administered 40 μL (split between each naris) of SARS-CoV-2 vaccine (5 μg or 25 μg) formulated in 2 different LNP compositions; as a control, 1 group (n = 10) was administered tris/sucrose buffer (mock-vaccination) intranasally. An additional 2 groups (n = 10 per group) were intramuscularly vaccinated into the hind leg with the SARS-CoV-2 vaccine (0.4 μg or 1 μg), formulated with the preclinical version of the same LNP utilized in mRNA-1273. Serum samples for immunogenicity assessments were collected at 3 weeks after dose 1 (Day 21) and 3 weeks after dose 2 (Day 41).

At 21 days after dose 2 (Day 42), all vaccinated hamsters were infected with 100 μL (50 μL/naris) SARS-CoV-2 (2019-nCoV/USA-WA1/2020; Genbank: MN985325.1; courtesy of World Reference Center for Emerging Viruses and Arboviruses, University of Texas Medical Branch) at 10^5^ PFU. Through 14 days after viral challenge, hamsters were monitored daily for weight changes. At 3 days and 14 days post-infection, lungs and nasal turbinates were collected from each vaccine group (n = 5 animals per timepoint). Prior to SARS-CoV-2 challenge, one animal each in the mRNA-LNP2 5 μg and 25 μg groups died: one succumbed to territorial behavior and the other cause of death was unknown. Animal experiments were carried out in compliance with approval from the Institutional Animal Care and Use Committee of the University of Texas Medical Branch.

### Preclinical mRNA and lipid nanoparticle production process

A sequence-optimized mRNA encoding the SARS-CoV-2 S protein with 6 proline mutations^32^ was in vitro synthesized and purified as previously described.^27^ mRNA was LNP-encapsulated via nanoprecipitation by microfluidic mixing of ionizable, structural, helper, and polyethylene glycol lipids in acetate buffer (pH 5.0), followed by buffer exchange, concentration via tangential flow filtration, and filtration through a 0.8/0.2 μm membrane;^27,48^ an additional lipid was added for mRNA-LNP2. The drug product was analytically characterized and the products were evaluated as acceptable for in vivo use.

### S-2P-specific ELISA

MaxiSorp 96-well flat-bottom plates (Thermo Fisher Scientific) were coated with 1 μg/mL (for IgG) or 5 μg/mL (for IgA) S-2P protein (GenScript), corresponding to the spike protein of the Wuhan-Hu-1 virus stabilized with 2 proline mutations, and incubated at 4°C overnight. The plates were then washed 4 times with PBS + 0.05% Tween-20 and blocked with SuperBlock buffer in PBS (Thermo Fisher Scientific) for 1.5 hours at 37°C. After washing, 5-fold serial dilutions of serum (assay diluent: PBS + 5% goat serum [Gibco] + 0.05% Tween-20) was added and incubated for 2 hours at 37°C (IgG) or overnight (IgA). Plates were washed and bound antibodies were detected with horseradish peroxidase (HRP)– conjugated goat anti-hamster IgG antibodies (1:10,000; Abcam; AB7146) or HRP-conjugated rabbit anti-hamster IgA antibodies (1:5,000; Brookwood Biomedical; sab3003) for 1 hour at 37°C. Plates were washed and bound antibody detected with SureBlue TMB substrate (Kirkegaard & Perry Labs, Inc.). After incubating at room temperature for 12 minutes, 3,3,5,5-tetramethylbenzidine stop solution (Kirkegaard & Perry Labs, Inc.) was added and absorbance was measured at 450 nM. GraphPad Prism (V 9.4.0) was used to determine titers using a 4-parameter logistic curve fit for IgG or defined as the reciprocal dilution at approximately optical density (OD) for IgA with baseline defined as 3-fold above the OD of the blank.

### SARS-CoV-2 neutralization assay

Two-fold dilutions of serum (heat inactivated, at an initial 1:10 dilution) were prepared in serum-free minimal essential media (MEM), then incubated with SARS-CoV-2 (2019-nCoV/USA-WA01/2020 at a final concentration of 100 PFU) at 37°C for 1 hour. Mixtures of virus-sera were then absorbed onto monolayers of Vero-E6 cells for 1 hour at 37°C in 96-well plates, then replaced with an overlay of MEM/methylcellulose/2% fetal bovine serum (FBS) and incubated for 2 days at 37°C in humidified 5% CO_2_. Plaques were immunostained as described below for viral load analysis by plaque assay and then counted with the ImmunoSpot analyzer (CTL); neutralization titers were determined at an endpoint of 60% plaque reduction.

### Analysis of viral load by plaque assay

Nasal turbinates and right lung were homogenized in Leibovitz L-15 medium (Thermo Fisher Scientific) supplemented with 10% FBS and 1x antibiotic-antimycotic by a TissueLyser II bead mill with 5-mm stainless steel beads (Qiagen). After brief centrifugation, 10-fold serial dilutions of homogenates were prepared in serum-free MEM, then absorbed on 48-well plates of Vero-E6 monolayers for 1 hour at 37°C. The virus inoculum was removed, replaced with an overlay of MEM/methylcellulose/2% FBS, and incubated for 3 days. Plaques were then immunostained using a human monoclonal antibody cocktail specific for the SARS-CoV-2 S protein (Clones DB_A03-09, 12; DB_B01-04, B07-10, 12; DB_C01-05, 07,09, 10; DB_D01, 02; DB_E01-04, 06, 07; DB_F02-03; kindly provided by Distributed Bio) and an anti-human IgG HRP-conjugated secondary antibody (Cat No. 5220-0456; Sera Care) and then counted to determine virus load per gram of tissue.

### Analysis of viral load by qRT-PCR

Replicating viral RNA in lung and nasal turbinates was determined via qRT-PCR measuring subgenomic SARS-CoV-2 E gene RNA using previously described primers, probe, and cycle conditions.^49^ In brief, RNA was extracted from homogenates using TRIzol LS (Thermo Fisher Scientific) and Direct-zol RNA Microprep kit (Zymo Research). Quantitative one-step real-time PCR was performed using extracted RNA (10 ng), TaqMan Fast Virus 1-step Master Mix (Thermo Fisher Scientific), primers, and a FAM-ZEN/Iowa Black FQ labeled probe sequence (Integrated DNA Technologies) on the QuantStudio 6 system (Applied Biosystems). An Ultramer DNA oligonucleotide spanning the amplicon (Integrated DNA Technologies) was used for standard curve generation to calculate subgenomic RNA copies per gram of tissue.

### Histopathology

Histological analysis of lung samples followed a standard protocol. In brief, the lower left lobe of the lung was fixed in 10% neutral buffered formalin, paraffin-embedded, sectioned (5 μm), and stained with hematoxylin and eosin (H&E). Sections were evaluated in a blinded manner by a board-certified veterinary pathologist under light microscopy with an Olympus BX51 microscope. Slides were scanned with a 20x (N.A. 0.8) objective at a single layer with continuous stage movement scanning method and images were captured using a Pannoramic 250 Flash III (3DHISTECH). Glass slides were examined, and microscopic diagnoses were graded independently on a 5-level severity scale (grades 1 to 5: minimal, mild, moderate, marked, and severe) by 2 veterinary pathologists.

### Immunohistochemistry

Immunohistochemistry (IHC) was performed on Formalin-Fixed Paraffin Embedded (FFPE) sections using the Lecia Bond RX auto-stainer (Lecia Microsystems). Sections were baked for one hour prior to staining and dewaxed on the instrument. Antigen retrieval was then performed for 20 minutes at 95°C using Lecia Epitope Retrieval Buffer 2 followed by treatment with Dako serum-free protein block (X090930-2, Agilent Dako) for 15 minutes to prevent non-specific binding of the antibody. Tissue was incubated with 0.083 μg/mL of anti-SARS-CoV-2 nucleocapsid protein (GTX135357, GeneTex) for 30 minutes and then detected using Bond Polymer Refine Detection Kit (DS9800, Lecia Microsystems) and bluing reagent (3802918, Lecia Microsystems) to enhance the color. Images were taken at 20x magnification using a Panoramic 250 Flash II scanner (3DHISTECH). Image analysis software was performed using Halo software (Indica Labs).

### Statistical modeling and hypothesis testing

Bayesian linear mixed model was used to model IgG, IgA and neutralization titers, separately. A Bayesian model was chosen for its flexibility in model estimation when the data was censored (left at the limit of detection) and presented heterogeneous group variances. Since the Bayesian model was employed for ease of model fitting, but not as a means to include prior information, we opt for non-informative prior in our analysis. For IgG, IgA and neutralization titers (log10 titers), each dosing day was modeled separately with one main effect of composition and dose combination (6 levels) and residual variance specific to each dose level (5 μg, 25 μg, 0.4 μg, and 1 μg). Default priors in the brms R package was used, with non-informative flat priors used for all regression coefficients. Holm’s method was used to adjust *P*-values for multiple comparisons. For viral loads (log10 transformed), ordinary linear regression was used with modeled data on Day 3 only as viral loads on Day 14 were zero for all hamsters. Sidak’s method was used to adjust *P*-values for multiple comparisons. All hypothesis testing was done two-sided at alpha level of 0.05, except when noted otherwise. R version 4.1.2 was used for statistical modelling.^50^

Kruskal-Wallis non-parametric test was implemented in hypothesis testing for image analysis using GraphPad’s Prism software. This form of ANOVA accounts for the small sample size in each experimental group, as well as the small percentage of N-protein positive cells among animals in the vaccinated groups.

## Data availability statement

The authors declare that the data supporting the findings of this study are available within this article and its Supplementary Information.

## Acknowledgments

We would like to thank the University of Texas Medical Branch Animal Resources Center for technical assistance, the World Reference Center for Emerging Viruses and Arboviruses at the University of Texas Medical Branch for generously providing the challenge virus. Medical writing and editorial assistance were provided by Emily Stackpole, PhD, and Wynand van Losenoord, MSc, of MEDiSTRAVA in accordance with Good Publication Practice (GPP3) guidelines, funded by Moderna, Inc., and under the direction of the authors.

## Author contributions

Study concept and design was performed by GBV, MM, AC, CEM, DEK, AB, and KB. Data collection was undertaken by GBV, MM, AG, AJ, CAP, CEM, MAH, MEC, JNW, AB, and KB. All authors contributed to the analysis/interpretation of the data, writing and/or review of the manuscript, and approved the final draft.

## References

1 World Health Organization. The top 10 causes of death, <https://www.who.int/news-room/fact-sheets/detail/the-top-10-causes-of-death> (2020).

2 World Health Organization. COVID-19 Vaccine Tracker and Landscape, <https://www.who.int/publications/m/item/draft-landscape-of-covid-19-candidate-vaccines> (2022).

3 Collaborators, G. B. D. L. R. I. Estimates of the global, regional, and national morbidity, mortality, and aetiologies of lower respiratory infections in 195 countries, 1990-2016: a systematic analysis for the Global Burden of Disease Study 2016. Lancet Infect Dis 18, 1191–1210, doi:10.1016/S1473-3099(18)30310-4 (2018).

4 McCloskey, B., Dar, O., Zumla, A. & Heymann, D. L. Emerging infectious diseases and pandemic potential: status quo and reducing risk of global spread. Lancet Infect Dis 14, 1001–1010, doi:10.1016/S1473-3099(14)70846-1 (2014).

5 Centers for Disease, C. & Prevention. *Ten great public health achievements--worldwide*, 2001-2010. MMWR Morb Mortal Wkly Rep 60, 814–818 (2011).

6 Lavelle, E. C. & Ward, R. W. Mucosal vaccines - fortifying the frontiers. Nat Rev Immunol 22, 236–250, doi:10.1038/s41577-021-00583-2 (2022).

7 Mouro, V. & Fischer, A. Dealing with a mucosal viral pandemic: lessons from COVID-19 vaccines. Mucosal Immunol 15, 584–594, doi:10.1038/s41385-022-00517-8 (2022).

8 Alu, A. et al. Intranasal COVID-19 vaccines: From bench to bed. EBioMedicine 76, 103841, doi:10.1016/j.ebiom.2022.103841 (2022).

9 Yusuf, H. & Kett, V. Current prospects and future challenges for nasal vaccine delivery. Hum Vaccin Immunother 13, 34–45, doi:10.1080/21645515.2016.1239668 (2017).

10 Hassan, A. O. et al. A Single-Dose Intranasal ChAd Vaccine Protects Upper and Lower Respiratory Tracts against SARS-CoV-2. Cell 183, 169–184.e113, doi:10.1016/j.cell.2020.08.026 (2020).

11 Krammer, F. SARS-CoV-2 vaccines in development. Nature 586, 516–527, doi:10.1038/s41586-020-2798-3 (2020).

12 Russell, M. W., Moldoveanu, Z., Ogra, P. L. & Mestecky, J. Mucosal Immunity in COVID-19: A Neglected but Critical Aspect of SARS-CoV-2 Infection. Front Immunol 11, 611337, doi:10.3389/fimmu.2020.611337 (2020).

13 Lapuente, D. et al. Protective mucosal immunity against SARS-CoV-2 after heterologous systemic prime-mucosal boost immunization. Nat Commun 12, 6871, doi:10.1038/s41467-021-27063-4 (2021).

14 Hartwell, B. L. et al. Intranasal vaccination with lipid-conjugated immunogens promotes antigen transmucosal uptake to drive mucosal and systemic immunity. Sci Transl Med 14, eabn1413, doi:10.1126/scitranslmed.abn1413 (2022).

15 van Doremalen, N. et al. Intranasal ChAdOx1 nCoV-19/AZD1222 vaccination reduces viral shedding after SARS-CoV-2 D614G challenge in preclinical models. Sci Transl Med 13, doi:10.1126/scitranslmed.abh0755 (2021).

16 Chavda, V. P., Vora, L. K., Pandya, A. K. & Patravale, V. B. Intranasal vaccines for SARS-CoV-2: From challenges to potential in COVID-19 management. Drug Discov Today 26, 2619–2636, doi:10.1016/j.drudis.2021.07.021 (2021).

17 Birkhoff, M., Leitz, M. & Marx, D. Advantages of Intranasal Vaccination and Considerations on Device Selection. Indian Journal of Pharmaceutical Sciences 71, 729–731 (2009).

18 Freeman, D. et al. Injection fears and COVID-19 vaccine hesitancy. Psychol Med, 1–11, doi:10.1017/S0033291721002609 (2021).

19 Waltz, E. China and India approve nasal COVID vaccines - are they a game changer? Nature 609, 450, doi:10.1038/d41586-022-02851-0 (2022).

20 Package Insert - SPIKEVAX, <https://www.fda.gov/media/155675/download> (2022).

21 Bruxvoort, K. et al. Real-World Effectiveness of the mRNA-1273 Vaccine Against COVID-19: Interim Results from a Prospective Observational Cohort Study. Preprints with THE LANCET, doi:Available at SSRN: https://ssrn.com/abstract=3916094 or http://dx.doi.org/10.2139/ssrn.3916094 (2021).

22 Chemaitelly, H. et al. mRNA-1273 COVID-19 vaccine effectiveness against the B.1.1.7 and B.1.351 variants and severe COVID-19 disease in Qatar. Nat Med,doi: 10.1038/s41591-021-01446-y (2021).

23 Dickerman, B. A. et al. Comparative Effectiveness of BNT162b2 and mRNA-1273 Vaccines in U.S. Veterans. New England Journal of Medicine 386, 105–115, doi:10.1056/NEJMoa2115463 (2021).

24 Polack, F. P. et al. Safety and efficacy of the BNT162b2 mRNA Covid-19 vaccine. N Engl J Med 383, 2603–2615, doi:10.1056/NEJMoa2034577 (2020).

25 Pilishvili, T. et al. Effectiveness of mRNA Covid-19 Vaccine among U.S. Health Care Personnel. N Engl J Med 385, e90, doi:10.1056/NEJMoa2106599 (2021).

26 Baden, L. R. et al. Efficacy and safety of the mRNA-1273 SARS-CoV-2 vaccine. N Engl J Med 384, 403–416, doi:10.1056/NEJMoa2035389 (2021).

27 Hassett, K. J. et al. Optimization of Lipid Nanoparticles for Intramuscular Administration of mRNA Vaccines. Mol Ther Nucleic Acids 15, 1–11, doi:10.1016/j.omtn.2019.01.013 (2019).

28 Mendonca, S. A., Lorincz, R., Boucher, P. & Curiel, D. T. Adenoviral vector vaccine platforms in the SARS-CoV-2 pandemic. NPJ Vaccines 6, 97, doi:10.1038/s41541-021-00356-x (2021).

29 Pardi, N., Hogan, M. J., Porter, F. W. & Weissman, D. mRNA vaccines - a new era in vaccinology. Nat Rev Drug Discov 17, 261–279, doi:10.1038/nrd.2017.243 (2018).

30 Veiga, N., Diesendruck, Y. & Peer, D. Targeted lipid nanoparticles for RNA therapeutics and immunomodulation in leukocytes. Adv Drug Deliv Rev 159, 364–376, doi:10.1016/j.addr.2020.04.002 (2020).

31 Tombacz, I. et al. Highly efficient CD4+ T cell targeting and genetic recombination using engineered CD4+ cell-homing mRNA-LNPs. Mol Ther 29, 3293–3304, doi:10.1016/j.ymthe.2021.06.004 (2021).

32 Hsieh, C. L. et al. Structure-based design of prefusion-stabilized SARS-CoV-2 spikes. Science 369, 1501–1505, doi:10.1126/science.abd0826 (2020).

33 Halfmann, P. J. et al. SARS-CoV-2 Omicron virus causes attenuated disease in mice and hamsters. Nature 603, 687–692, doi:10.1038/s41586-022-04441-6 (2022).

34 McMahan, K. et al. Reduced pathogenicity of the SARS-CoV-2 omicron variant in hamsters. Med (N Y) 3, 262–268 e264, doi:10.1016/j.medj.2022.03.004 (2022).

35 Meyer, M. et al. Attenuated activation of pulmonary immune cells in mRNA-1273-vaccinated hamsters after SARS-CoV-2 infection. J Clin Invest 131, doi:10.1172/JCI148036 (2021).

36 Imai, M. et al. Syrian hamsters as a small animal model for SARS-CoV-2 infection and countermeasure development. Proc Natl Acad Sci U S A 117, 16587–16595, doi:10.1073/pnas.2009799117 (2020).

37 Bharat Biotech International Limited. iNCOVACC®, World’s first Intranasal Vaccine to receive both Primary series and Heterologous booster approval, <https://www.bharatbiotech.com/images/press/bharat-biotech-incovacc-booster-approval-press-release.pdf> (2022).

38 CanSino Biologics Inc. CanSinoBIO’s Convidecia Air™ Receives Approval in China, <https://www.cansinotech.com/html/1/179/180/1100.html> (2022).

39 Wong, T. Y. et al. Intranasal administration of BReC-CoV-2 COVID-19 vaccine protects K18-hACE2 mice against lethal SARS-CoV-2 challenge. npj Vaccines 7, 36, doi:10.1038/s41541-022-00451-7 (2022).

40 Zhang, Z. et al. Aerosolized Ad5-nCoV booster vaccination elicited potent immune response against the SARS-CoV-2 Omicron variant after inactivated COVID-19 vaccine priming. medRxiv, 2022.2003.2008.22271816, doi:10.1101/2022.03.08.22271816 (2022).

41 Tioni, M. F. et al. One mucosal administration of a live attenuated recombinant COVID-19 vaccine protects nonhuman primates from SARS-CoV-2. bioRxiv, 2021.2007.2016.452733, doi:10.1101/2021.07.16.452733 (2021).

42 Madhavan, M. et al. Tolerability and immunogenicity of an intranasally-administered adenovirus-vectored COVID-19 vaccine: An open-label partially-randomised ascending dose phase I trial. EBioMedicine, 104298, doi:10.1016/j.ebiom.2022.104298 (2022).

43 Edwards, D. K. & Carfi, A. Messenger ribonucleic acid vaccines against infectious diseases: current concepts and future prospects. Curr Opin Immunol 77, 102214, doi:10.1016/j.coi.2022.102214 (2022).

44 Choi, A. et al. Safety and immunogenicity of SARS-CoV-2 variant mRNA vaccine boosters in healthy adults: an interim analysis. Nat Med, doi:10.1038/s41591-021-01527-y (2021).

45 Bruxvoort, K. J. et al. Real-world effectiveness of the mRNA-1273 vaccine against COVID-19: Interim results from a prospective observational cohort study. Lancet Reg Health Am, 100134, doi:10.1016/j.lana.2021.100134 (2021).

46 El Sahly, H. M. et al. Efficacy of the mRNA-1273 SARS-CoV-2 Vaccine at Completion of Blinded Phase. N Engl J Med 385, 1774–1785, doi:10.1056/NEJMoa2113017 (2021).

47 Chan, J. F. et al. Simulation of the Clinical and Pathological Manifestations of Coronavirus Disease 2019 (COVID-19) in a Golden Syrian Hamster Model: Implications for Disease Pathogenesis and Transmissibility. Clin Infect Dis 71, 2428–2446, doi:10.1093/cid/ciaa325 (2020).

48 Sabnis, S. et al. A Novel Amino Lipid Series for mRNA Delivery: Improved Endosomal Escape and Sustained Pharmacology and Safety in Non-human Primates. Mol Ther 26, 1509–1519, doi:10.1016/j.ymthe.2018.03.010 (2018).

49 Wolfel, R. et al. Virological assessment of hospitalized patients with COVID-2019. Nature 581, 465–469, doi:10.1038/s41586-020-2196-x (2020).

50 R Core Team. R: A language and environment for statistical computing. R Foundation for Statistical Computing, Vienna, Austria. URL https://www.R-project.org/.2021).

