## Supplemental for "Intranasal mRNA-LNP vaccination protects hamsters from SARS-CoV-2 infection"

### Supplemental information

***Supplementary Table 1. Statistical comparisons of S-specific serum binding IgG antibody titers after vaccination***

| Comparison | Estimated Fold-change (95% CI) | Adjusted P-value |
|---|---|---|
| *Dose 1 (Day 21)* | | |
| 25-µg mRNA-LNP1 over 5-µg mRNA-LNP1 | 68.53 (25.63-185.69) | <1e-6 |
| 25-µg mRNA-LNP2 over 5-µg mRNA-LNP1 | 67.68 (25.04-182.43) | <1e-6 |
| 5-µg mRNA-LNP2 over 5-µg mRNA-LNP1 | 50.61 (22.1-117.24) | <1e-6 |
| 25-µg mRNA-LNP1 over 0.4-µg IM | 3.11 (0.86-11.06) | 0.9619 |
| 25-µg mRNA-LNP1 over 1-µg IM | 1.63 (0.61-4.38) | 0.8426 |
| 5-µg mRNA-LNP1 over 0.4-µg IM | 0.05 (0.02-0.14) | <1e-6 |
| 5-µg mRNA-LNP1 over 1-µg IM | 0.02 (0.01-0.05) | <1e-6 |
| 25-µg mRNA-LNP2 over 25-µg mRNA-LNP1 | 0.99 (0.31-3.14) | 0.4892 |
| 25-µg mRNA-LNP2 over 5-µg mRNA-LNP2 | 1.34 (0.5-3.71) | 0.7183 |
| 25-µg mRNA-LNP2 over 0.4-µg IM | 3.07 (0.9-11.02) | 0.9635 |
| 25-µg mRNA-LNP2 over 1-µg IM | 1.61 (0.6-4.4) | 0.8312 |
| 5-µg mRNA-LNP2 over 25-µg mRNA-LNP1 | 0.74 (0.27-2.03) | 0.2668 |
| 5-µg mRNA-LNP2 over 0.4-µg IM | 2.3 (0.75-6.89) | 0.9359 |
| 5-µg mRNA-LNP2 over 1-µg IM | 1.2 (0.54-2.69) | 0.6883 |
| 1-µg IM over 0.4-µg IM | 1.91 (0.63-5.83) | 0.8853 |
| *Dose 2 (Day 41)* | | |
| 25-µg mRNA-LNP1 over 5-µg mRNA-LNP1 | 3.64 (1.54-8.55) | 0.0024 |
| 25-µg mRNA-LNP2 over 25-µg mRNA-LNP1 | 3.68 (1.78-7.55) | 6e-04 |
| 25-µg mRNA-LNP2 over 5-µg mRNA-LNP1 | 13.38 (5.58-31.97) | <1e-6 |
| 25-µg mRNA-LNP2 over 5-µg mRNA-LNP2 | 2.7 (1.1-6.61) | 0.0156 |
| 5-µg mRNA-LNP2 over 5-µg mRNA-LNP1 | 4.95 (1.8-13.8) | 0.0023 |
| 25-µg mRNA-LNP1 over 0.4-µg IM | 0.61 (0.27-1.34) | 0.1004 |
| 25-µg mRNA-LNP1 over 1-µg IM | 0.37 (0.17-0.86) | 0.0138 |
| 5-µg mRNA-LNP1 over 0.4-µg IM | 0.17 (0.06-0.43) | 2e-04 |
| 5-µg mRNA-LNP1 over 1-µg IM | 0.1 (0.04-0.27) | 2e-04 |

| | | |
|---|---|---|
| 25-µg mRNA-LNP2 over 0.4-µg IM | 2.24 (0.99-4.93) | 0.9742 |
| 25-µg mRNA-LNP2 over 1-µg IM | 1.38 (0.61-3.2) | 0.7796 |
| 5-µg mRNA-LNP2 over 25-µg mRNA-LNP1 | 1.36 (0.56-3.32) | 0.7592 |
| 5-µg mRNA-LNP2 over 0.4-µg IM | 0.83 (0.32-2.27) | 0.3406 |
| 5-µg mRNA-LNP2 over 1-µg IM | 0.51 (0.19-1.38) | 0.0894 |
| 1-µg IM over 0.4-µg IM | 1.63 (0.63-3.98) | 0.8621 |

CI, confidence interval; IgG, immunoglobulin G; IM, intramuscular; LNP, lipid nanoparticle; mRNA, messenger RNA.

***Supplementary Table 2. Statistical comparisons of S-specific serum binding IgA antibody titers after vaccination***

| **Comparison** | **Estimated Fold-change (95% CI)** | **Adjusted P-value** |
|---|---|---|
| *Dose 1 (Day 21)* | | |
| 25-µg mRNA-LNP2 over 25-µg mRNA-LNP1 | 83.81 (38.04-224.25) | <1e-6 |
| 25-µg mRNA-LNP2 over 5-µg mRNA-LNP2 | 32.7 (3.92-716.22) | 0.0035 |
| 25-µg mRNA-LNP2 over 0.4-µg IM | 2.84 (1.18-6.73) | 0.0104 |
| 25-µg mRNA-LNP1 over 0.4-µg IM | 0.03 (0.01-0.09) | <1e-6 |
| 25-µg mRNA-LNP1 over 1-µg IM | 0.02 (0.01-0.05) | <1e-6 |
| 25-µg mRNA-LNP2 over 1-µg IM | 1.7 (0.72-3.93) | 0.8993 |
| 5-µg mRNA-LNP2 over 25-µg mRNA-LNP1 | 2.56 (0.12-24.18) | 0.8043 |
| 5-µg mRNA-LNP2 over 0.4-µg IM | 0.09 (<1e-6-0.8) | 0.0171 |
| 5-µg mRNA-LNP2 over 1-µg IM | 0.05 (<1e-6-0.44) | 0.0066 |
| 1-µg IM over 0.4-µg IM | 1.67 (0.67-4.14) | 0.8746 |
| *Dose 2 (Day 41)* | | |
| 25-µg mRNA-LNP1 over 5-µg mRNA-LNP1 | 3.74 (1.37-10.19) | 0.0051 |
| 25-µg mRNA-LNP2 over 25-µg mRNA-LNP1 | 5.77 (2.57-12.95) | <1e-6 |
| 25-µg mRNA-LNP2 over 5-µg mRNA-LNP1 | 21.57 (7.93-60.5) | <1e-6 |
| 25-µg mRNA-LNP2 over 5-µg mRNA-LNP2 | 4.05 (1.37-11.73) | 0.0066 |
| 5-µg mRNA-LNP2 over 5-µg mRNA-LNP1 | 5.33 (1.55-18.18) | 0.006 |
| 25-µg mRNA-LNP1 over 0.4-µg IM | 0.21 (0.08-0.52) | 0.0025 |
| 25-µg mRNA-LNP1 over 1-µg IM | 0.13 (0.05-0.33) | <1e-6 |
| 5-µg mRNA-LNP1 over 0.4-µg IM | 0.06 (0.02-0.18) | <1e-6 |
| 5-µg mRNA-LNP1 over 1-µg IM | 0.03 (0.01-0.1) | <1e-6 |
| 25-µg mRNA-LNP2 over 0.4-µg IM | 1.21 (0.47-3.09) | 0.6658 |
| 25-µg mRNA-LNP2 over 1-µg IM | 0.73 (0.28-1.88) | 0.2411 |
| 5-µg mRNA-LNP2 over 25-µg mRNA-LNP1 | 1.43 (0.51-4.12) | 0.7531 |
| 5-µg mRNA-LNP2 over 0.4-µg IM | 0.3 (0.09-0.95) | 0.0219 |
| 5-µg mRNA-LNP2 over 1-µg IM | 0.18 (0.06-0.57) | 0.0044 |
| 1-µg IM over 0.4-µg IM | 1.67 (0.58-4.79) | 0.8428 |

CI, confidence interval; IgA, immunoglobulin A; IM, intramuscular; LNP, lipid nanoparticle; mRNA, messenger RNA.

***Supplementary Table 3. Statistical comparisons of serum neutralizing antibody titers after vaccination***

| Comparison | Estimated Fold-change (95% CI) | Adjusted P-value |
|---|---|---|
| *Dose 1 (Day 21)* | | |
| 25-µg mRNA-LNP2 over 25-µg mRNA-LNP1 | 38.84 (8.7-287.37) | <1e-6 |
| 25-µg mRNA-LNP2 over 5-µg mRNA-LNP2 | 17.46 (2.21-275.24) | 0.0044 |
| 25-µg mRNA-LNP2 over 0.4-µg IM | 8.09 (1.09-96.32) | 0.0205 |
| 25-µg mRNA-LNP1 over 0.4-µg IM | 0.21 (0.02-2.64) | 0.0801 |
| 25-µg mRNA-LNP1 over 1-µg IM | 0.06 (0.01-0.21) | <1e-6 |
| 25-µg mRNA-LNP2 over 1-µg IM | 2.16 (0.63-7.5) | 0.8996 |
| 5-µg mRNA-LNP2 over 25-µg mRNA-LNP1 | 2.22 (0.12-24.4) | 0.7781 |
| 5-µg mRNA-LNP2 over 0.4-µg IM | 0.46 (0.02-7.72) | 0.275 |
| 5-µg mRNA-LNP2 over 1-µg IM | 0.12 (0.01-0.77) | 0.0149 |
| 1-µg IM over 0.4-µg IM | 3.74 (0.6-39.29) | 0.9309 |
| *Dose 2 (Day 41)* | | |
| 25-µg mRNA-LNP1 over 5-µg mRNA-LNP1 | 5.07 (1.53-18.22) | 0.0054 |
| 25-µg mRNA-LNP2 over 25-µg mRNA-LNP1 | 5.7 (2.28-14.07) | 0.0008 |
| 25-µg mRNA-LNP2 over 5-µg mRNA-LNP1 | 28.89 (8.5-111.28) | <1e-6 |
| 25-µg mRNA-LNP2 over 5-µg mRNA-LNP2 | 5.08 (1.46-18.06) | 0.0059 |
| 5-µg mRNA-LNP2 over 5-µg mRNA-LNP1 | 5.69 (1.32-27.13) | 0.0119 |
| 25-µg mRNA-LNP1 over 0.4-µg IM | 0.36 (0.14-0.91) | 0.0173 |
| 25-µg mRNA-LNP1 over 1-µg IM | 0.17 (0.06-0.45) | 0.0004 |
| 5-µg mRNA-LNP1 over 0.4-µg IM | 0.07 (0.02-.024) | 0.0002 |
| 5-µg mRNA-LNP1 over 1-µg IM | 0.03 (0.01-0.12) | <1e-6 |
| 25-µg mRNA-LNP2 over 0.4-µg IM | 2.06 (0.83-5.26) | 0.94 |
| 25-µg mRNA-LNP2 over 1-µg IM | 0.96 (0.33-2.65) | 0.4737 |
| 5-µg mRNA-LNP2 over 25-µg mRNA-LNP1 | 1.12 (0.32-3.86) | 0.5776 |
| 5-µg mRNA-LNP2 over 0.4-µg IM | 0.41 (0.12-1.43) | 0.0758 |
| 5-µg mRNA-LNP2 over 1-µg IM | 0.19 (0.05-0.69) | 0.0065 |
| 1-µg IM over 0.4-µg IM | 2.15 (0.77-6.24) | 0.9333 |

CI, confidence interval, IM, intramuscular; LNP, lipid nanoparticle; mRNA, messenger RNA.

***Supplementary Table 4. Statistical comparisons of viral load by plaque assay[a] in the lungs of vaccinated hamsters at 3 days after SARS-CoV-2 challenge***

| Comparison | Fold-change (standard error) | *t*-ratio | P-value |
|---|---|---|---|
| 0.4-µg IM over 1-µg IM | 2.32 (1.23) | 1.88 | 0.783 |
| 0.4-µg IM over 25-µg mRNA-LNP1 | 1.15 (1.17) | 0.98 | 1 |
| 0.4-µg IM over 5-µg mRNA-LNP1 | -1.91 (1.11) | -1.72 | 0.881 |
| 0.4-µg IM over 25-µg mRNA-LNP2 | 2.77 (1.05) | 2.64 | 0.245 |
| 0.4-µg IM over 5-µg mRNA-LNP2 | 0.99 (0.98) | 1.01 | 1 |
| 0.4-µg IM over mock | -2.60 (0.90) | -2.88 | 0.148 |
| 1-µg IM over 25-µg mRNA-LNP1 | 0.01 (1.23) | 0.01 | 1 |
| 1-µg IM over 5-µg mRNA-LNP1 | -3.05 (1.17) | -2.60 | 0.266 |
| 1-µg IM over 25-µg mRNA-LNP2 | 1.63 (1.11) | 1.46 | 0.97 |
| 1-µg IM over 5-µg mRNA-LNP2 | -0.15 (1.05) | -0.15 | 1 |
| 1-µg IM over mock | -3.73 (0.98) | -3.82 | 0.014 |
| 25-µg mRNA-LNP1 over 5-µg mRNA-LNP1 | -1.88 (1.23) | -1.53 | 0.955 |
| 25-µg mRNA-LNP1 over 25-µg mRNA-LNP2 | 2.79 (1.17) | 2.39 | 0.401 |
| 25-µg mRNA-LNP1 over 5-µg mRNA-LNP2 | 1.02 (1.11) | 0.91 | 1 |
| 25-µg mRNA-LNP1 over mock | -2.57 (1.05) | -2.45 | 0.354 |
| 5-µg mRNA-LNP1 over 25-µg mRNA-LNP2 | 5.85 (1.23) | 4.76 | 0.001 |
| 5-µg mRNA-LNP1 over 5-µg mRNA-LNP2 | 4.07 (1.17) | 3.48 | 0.035 |
| 5-µg mRNA-LNP1 over mock | 0.49 (1.11) | 0.44 | 1 |
| 25-µg mRNA-LNP2 over 5-µg mRNA-LNP2 | -0.60 (1.23) | -0.49 | 1 |
| 25-µg mRNA-LNP2 over mock | -4.18 (1.17) | -3.57 | 0.027 |
| 5-µg mRNA-LNP2 over mock | -2.40 (1.23) | -1.96 | 0.731 |

[a]Viral loads ($log_{10}$ transformed) were assessed by ordinary linear regression; only modeled data at Day 3 after challenge was evaluated since viral loads on Day 14 were zero for all hamsters. A degree of freedom of 28 was used.

IM, intramuscular; LNP, lipid nanoparticle; mRNA, messenger RNA.

***Supplementary Table 5. Statistical comparisons of viral load by plaque assay in the nasal turbinates of vaccinated hamsters at 3 days after SARS-CoV-2 challenge***

| Comparison | Fold-change (standard error) | *t*-ratio | P-value |
|---|---|---|---|
| 0.4-µg IM over 1-µg IM | 3.09 (0.87) | 3.56 | 0.028 |
| 0.4-µg IM over 25-µg mRNA-LNP1 | 0.55 (0.83) | 0.67 | 1 |
| 0.4-µg IM over 5-µg mRNA-LNP1 | -0.97 (0.78) | -1.24 | 0.995 |
| 0.4-µg IM over 25-µg mRNA-LNP2 | 2.59 (0.74) | 3.52 | 0.031 |
| 0.4-µg IM over 5-µg mRNA-LNP2 | 1.30 (0.69) | 1.89 | 0.779 |
| 0.4-µg IM over mock | -2.02 (0.64) | -3.18 | 0.073 |
| 1-µg IM over 25-µg mRNA-LNP1 | -1.53 (0.87) | -1.76 | 0.86 |
| 1-µg IM over 5-µg mRNA-LNP1 | -3.05 (0.83) | -3.69 | 0.02 |
| 1-µg IM over 25-µg mRNA-LNP2 | 0.52 (0.78) | 0.66 | 1 |
| 1-µg IM over 5-µg mRNA-LNP2 | -0.78 (0.74) | -1.05 | 0.999 |
| 1-µg IM over mock | -4.09 (0.69) | -5.95 | 0 |
| 25-µg mRNA-LNP1 over 5-µg mRNA-LNP1 | -0.51 (0.87) | -0.59 | 1 |
| 25-µg mRNA-LNP1 over 25-µg mRNA-LNP2 | 3.06 (0.83) | 3.70 | 0.019 |
| 25-µg mRNA-LNP1 over 5-µg mRNA-LNP2 | 1.76 (0.78) | 2.25 | 0.5 |
| 25-µg mRNA-LNP1 over mock | -1.56 (0.74) | -2.11 | 0.607 |
| 5-µg mRNA-LNP1 over 25-µg mRNA-LNP2 | 4.58 (0.87) | 5.29 | 0 |
| 5-µg mRNA-LNP1 over 5-µg mRNA-LNP2 | 3.28 (0.83) | 3.98 | 0.009 |
| 5-µg mRNA-LNP1 over mock | -0.03 (0.78) | -0.04 | 1 |
| 25-µg mRNA-LNP2 over 5-µg mRNA-LNP2 | -0.28 (0.87) | -0.33 | 1 |
| 25-µg mRNA-LNP2 over mock | -3.60 (0.83) | -4.36 | 0.003 |
| 5-µg mRNA-LNP2 over mock | -2.31 (0.87) | -2.66 | 0.236 |

[a]Viral loads ($log_{10}$ transformed) were assessed by ordinary linear regression; only modeled data at Day 3 after challenge was evaluated since viral loads on Day 14 were zero for all hamsters. A degree of freedom of 28 was used.

IM, intramuscular; LNP, lipid nanoparticle; mRNA, messenger RNA.

***Supplementary Table 6. Major pulmonary histopathological findings and severity scores after SARS-CoV-2 challenge by vaccine group.***

| ***Day 3 (After Challenge)*** | | | | | | | |
|---|---|---|---|---|---|---|---|
| | | **Intranasal** | | | | **Intramuscular** | |
| | Mock (n = 5) | mRNA-LNP1 5 µg (n = 5) | mRNA-LNP1 25 µg (n = 5) | mRNA-LNP2 5 µg (n = 5) | mRNA-LNP2 25 µg (n = 5) | 0.4 µg (n = 5) | 1 µg (n = 5) |
| Interstitial inflammation, n | | | | | | | |
| Group total | 5 | 5 | 5 | 5 | 5 | 5 | 5 |
| 2 (mild) | 2 | 4 | 2 | 2 | 1 | 5 | 2 |
| 3 (moderate) | 3 | 1 | 3 | 3 | 4 | 0 | 3 |
| Bronchial/bronchiolar inflammation, n | | | | | | | |
| Group total | 4 | 5 | 1 | 3 | 1 | 3 | 4 |
| 1 (minimal) | 2 | 1 | 1 | 1 | 0 | 2 | 3 |
| 2 (mild) | 0 | 3 | 0 | 2 | 1 | 1 | 1 |
| 3 (moderate) | 2 | 1 | 0 | 0 | 0 | 0 | 0 |
| Vascular inflammation, n | | | | | | | |
| Group total | 4 | 3 | 2 | 3 | 2 | 4 | 1 |
| 1 (minimal) | 0 | 1 | 1 | 1 | 1 | 3 | 0 |
| 2 (mild) | 2 | 2 | 1 | 1 | 0 | 1 | 1 |
| 3 (moderate) | 2 | 0 | 0 | 1 | 1 | 0 | 0 |
| ***Day 14 (After Challenge)*** | | | | | | | |
| | | **Intranasal** | | | | **Intramuscular** | |
| | Mock (n = 5) | mRNA-LNP1 5 µg (n = 5) | mRNA-LNP1 25 µg (n = 5) | mRNA-LNP2 5 µg (n = 4) | mRNA-LNP2 25 µg (n = 4) | 0.4 µg (n = 5) | 1 µg (n = 5) |
| Interstitial inflammation, n | | | | | | | |
| Group total | 5 | 5 | 5 | 4 | 4 | 5 | 5 |
| 2 (mild) | 0 | 5 | 4 | 0 | 4 | 4 | 3 |
| 3 (moderate) | 5 | 0 | 1 | 4 | 0 | 1 | 2 |
| Type II pneumocyte hyperplasia, n | | | | | | | |
| Group total | 5 | 4 | 5 | 4 | 2 | 1 | 5 |
| 1 (minimal) | 0 | 3 | 3 | 1 | 2 | 0 | 0 |
| 2 (mild) | 4 | 1 | 1 | 2 | 0 | 0 | 3 |
| 3 (moderate) | 1 | 0 | 1 | 1 | 0 | 1 | 2 |
| Bronchial/bronchiolar inflammation, n | | | | | | | |
| Group total | 4 | 3 | 2 | 3 | 1 | 1 | 3 |
| 1 (minimal) | 2 | 3 | 2 | 1 | 1 | 1 | 3 |
| 2 (mild) | 2 | 0 | 0 | 2 | 0 | 0 | 0 |
| Vascular inflammation, n | | | | | | | |
| Group total | 1 | 4 | 4 | 4 | 1 | 5 | 3 |
| 1 (minimal) | 1 | 4 | 4 | 4 | 1 | 5 | 3 |

IM, intramuscular; LNP, lipid nanoparticle; mRNA, messenger RNA; SARS-CoV-2, severe acute respiratory syndrome coronavirus 2.

***Supplementary Table 7. Summary of hamsters positive for SARS-CoV-2 N-protein at 3 days after challenge***

| **Vaccine Group** | **Administration Route** | **Ratio of N-protein Positive Hamsters to Total Population (n/N)** |
|---|---|---|
| Mock | IN | 5/5 |
| mRNA-LNP1 5 µg | IN | 5/5 |
| mRNA-LNP1 25 µg | IN | 2/5 |
| mRNA-LNP2 5 µg | IN | 4/5 |
| mRNA-LNP2 25 µg | IN | 4/5 |
| Intramuscular 0.4 µg | IM | 3/5 |
| Intramuscular 1 µg | IM | 2/5 |

IN, intranasal; IM, intramuscular; LNP, lipid nanoparticle; mRNA, messenger RNA; N protein, nucleocapsid protein; SARS-CoV-2, severe acute respiratory syndrome coronavirus 2.

***Supplementary Figure 1. Viral load as determined via qRT-PCR through 14 days after SARS-CoV-2 challenge in vaccinated hamsters.***

Viral load (sgRNA copies per gram of tissue) at 3 days and 14 days after SARS-CoV-2 challenge in (a) lungs and (b) nasal turbinates of vaccinated hamsters. Animal-level data are shown as dots (n = 5 animals per group), with the grey lines representing the geometric mean of each group. LLOD = $10^3$ copies/g of tissue.

IM, intramuscular; IN, intranasal; LNP, lipid nanoparticle; mRNA, messenger RNA; qRT-PCR, quantitative reverse transcription polymerase chain reaction; SARS-CoV-2, severe acute respiratory syndrome coronavirus 2; sgRNA, subgenomic RNA.

**a**

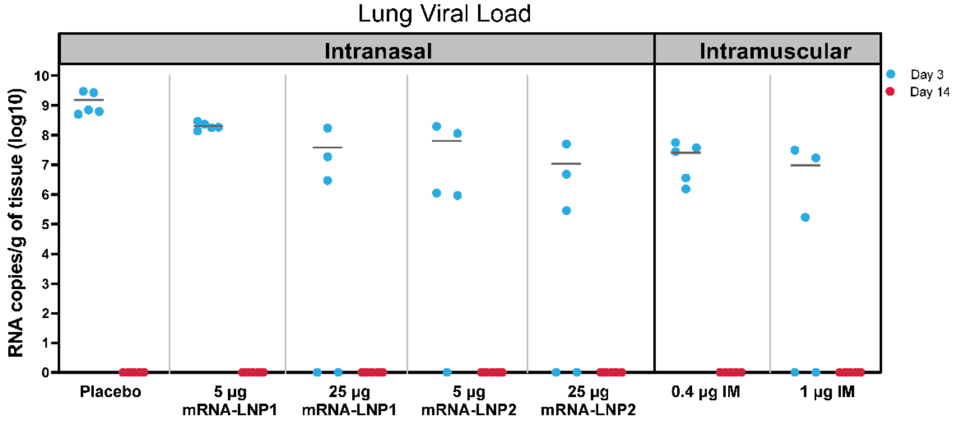


**b**

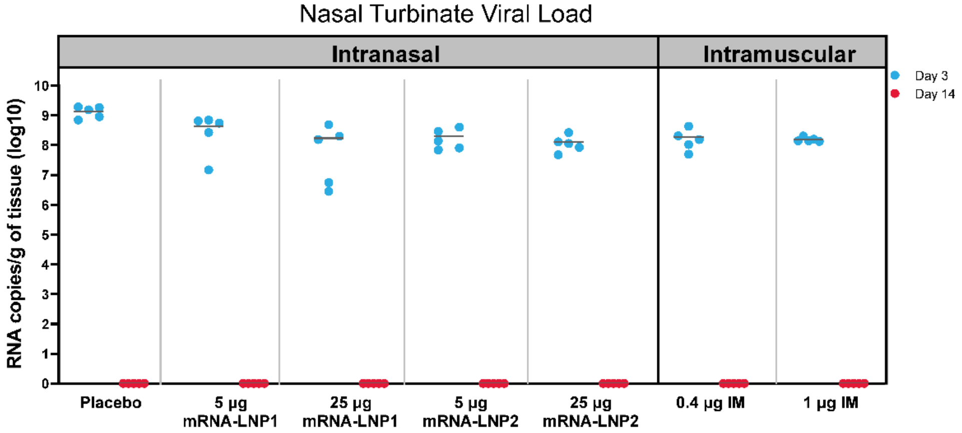

***Supplementary Figure 2. Pulmonary pathology characteristics at 14 days after SARS-CoV-2 challenge in vaccinated hamsters.***

Lung sections from hamsters at 14 days after SARS-CoV-2 challenge were stained with H&E. Representative images of (a) interstitial inflammation, (b) type II pneumocyte hyperplasia (arrows), or (c) airways and blood vessels are shown for hamsters intranasally administered 2 doses of tris/sucrose buffer (mock-vaccinated), mRNA-LNP1 (25 µg), mRNA-LNP2 (25 µg), or were intramuscularly vaccinated with 2 doses of vaccine (1.0 µg). Scale bars = 100 µm.

H&E, hematoxylin and eosin; IN, intranasal; SARS-CoV-2, severe acute respiratory syndrome coronavirus 2.

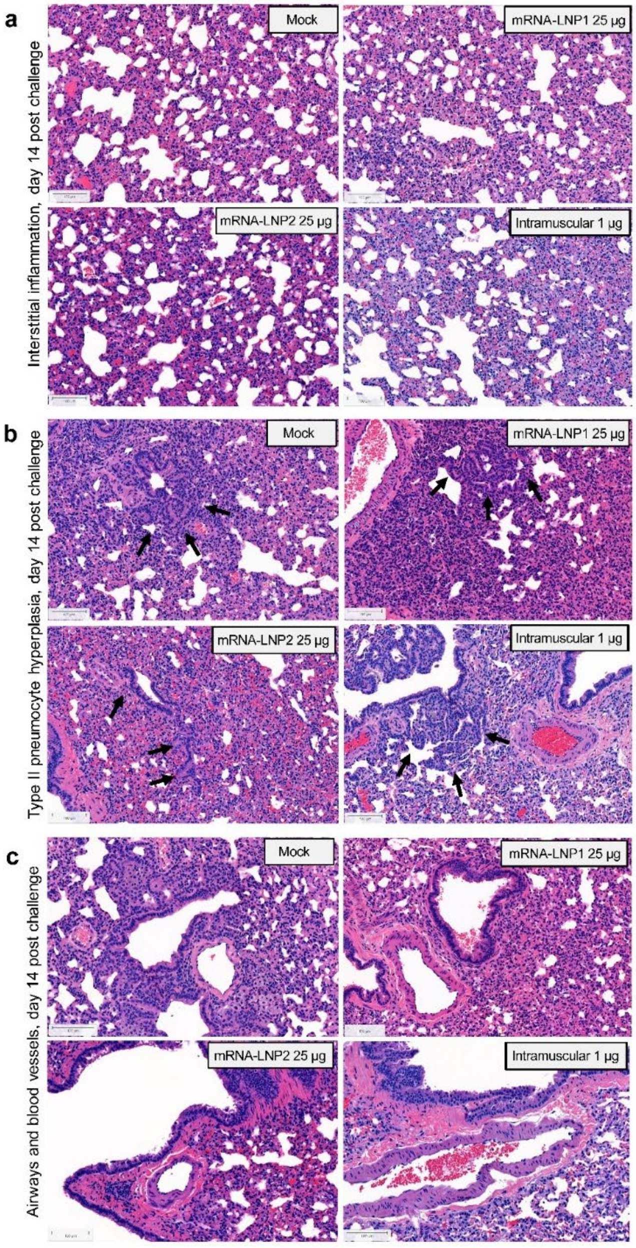
